# Macronutrient balance differentially regulates growth, immune function, and oxidative stress responses in *Plutella xylostella*

**DOI:** 10.64898/2026.07.27.740871

**Authors:** Anwesha Ghosh, Radhika Venkatesan

## Abstract

Nutrition plays a fundamental role in insect physiology, yet how variation in dietary macronutrient balance shapes multiple physiological functions remains poorly understood. Using the Geometric Framework for Nutrition, we examined how dietary protein and carbohydrate composition influence growth, immunity and stress physiology in the specialist herbivore *Plutella xylostella*. Growth depended jointly on protein and carbohydrate availability, whereas immune and oxidative stress traits showed distinct response surfaces across the tested nutrient landscape. Total hemocyte counts and phenoloxidase activity were highest at intermediate protein levels, and survival following *Bacillus thuringiensis* challenge was likewise associated primarily with protein availability, pointing to a shared nutritional basis for immunity and pathogen resistance. Antioxidant and detoxification enzymes showed trait-specific nutrient responses, indicating that different physiological systems are favored by different macronutrient combinations. No single protein-carbohydrate ratio within the tested landscape simultaneously supported growth, immune performance, and oxidative homeostasis, highlighting trade-offs associated with nutrient allocation across physiological systems. These findings advance understanding of how nutrient-dependent mechanisms coordinate – and constrain – multiple physiological functions in a specialist herbivore.

## 1. Introduction

Resource availability fundamentally shapes organismal performance and life-history strategies. In holometabolous insects, the larval stage is particularly critical, as resource acquisition and allocation during this period determine immediate growth and survival while also influencing the adult phenotype after metamorphosis. Variation in both the quantity and quality of nutrients obtained during larval development can generate carry-over effects that extend into pupation and adulthood, ultimately impacting subsequent life-history traits (Dmitriew & Rowe, 2011; Niitepõld & Boggs, 2022). Larvae must also face the challenge of balancing somatic growth with immune defense against pathogens, as immune activation demands considerable metabolic resources and can compete with other fitness-related processes for limited nutrients (Ponton *et al*., 2011; Schwenke *et al*., 2016). Life-history theory predicts that such competing demands generate intrinsic trade-offs, since investment in one function may carry costs for others (Stearns, 1989). In herbivorous insects, these competing demands make the balance of dietary macronutrients - particularly protein (P) and carbohydrate (C) - a central factor in determining how resources are allocated among growth, defense, and physiological maintenance (Lee et *al*., 2008; Povey *et al*., 2009; Cotter *et al*., 2011; Kazek *et al*., 2024). The Geometric Framework for Nutrition (GFN) offers a robust approach for quantifying how macronutrient balance modulates physiological responses and for identifying the nutritional conditions associated with specific outcomes (Simpson & Raubenheimer, 1999). Whether multiple physiological functions converge on shared nutritional conditions or are subject to distinct constraints across the nutritional landscape remains less understood - particularly in specialist herbivores, where this gap limits our understanding of how dietary variation translates into coordinated or conflicting trait responses under ecologically relevant nutritional conditions.

Previous studies of macronutrient balance and performance have focused predominantly on generalist herbivores with broad dietary breadths in which larvae can compensate for suboptimal macronutrient intake by switching among hosts (Lee *et al*., 2002; Cotter *et al*., 2011; Deans *et al*., 2015; Cotter *et al*., 2019). Specialist herbivores, by contrast, function within a relatively constrained nutritional space defined by their limited host-plant range, despite having evolved various adaptations to tolerate or circumvent host plant defenses (Diamond & Kingsolver, 2011; Ali & Agrawal, 2012). This constraint may intensify trade-offs among growth, immune function, and stress tolerance (Wilson *et al*., 2019), making specialist herbivores particularly informative systems for investigating how nutrient balance shapes physiological performance.

Immune responses in insects are primarily innate, relying on hemocyte activity and enzymatic pathways such as the prophenoloxidase cascade leading to melanization. The melanization reaction generates reactive quinones and other reactive by-products that contribute to antimicrobial defense while also imposing an oxidative burden on host tissues thereby linking immune activation to oxidative regulation (González□Santoyo & Córdoba-Aguilar, 2012; Zhao *et al*., 2007). Beyond immune activation, these insects encounter reactive oxygen species generated through aerobic metabolism during the processing of dietary constituents (Barbehenn, 2002; Felton & Summers, 1995). The insect midgut possesses a diverse antioxidant defense system, including superoxide dismutases, catalases, and peroxidases, which collectively scavenge reactive oxygen species and contribute to redox homeostasis (Barbehenn, 2002; Xia *et al*., 2017; Chamani *et al*., 2025). Detoxification represents another physiologically demanding process in herbivorous insects, as the biotransformation and elimination of dietary and host-derived compounds require coordinated activity of enzyme systems such as glutathione S-transferases and esterases; because the biosynthesis of these enzymes carry energetic costs, detoxification may impose additional nutritional demands that reduce resources available for other fitness-related functions (Castañeda *et al*., 2009).

Previous work has shown that individual physiological traits can respond differently to dietary protein and carbohydrate intake (Ponton *et al*., 2011; Wilson *et al*., 2019), suggesting that distinct traits may peak under different macronutrient conditions. If so, no single nutritional state may simultaneously favor growth, defense, and stress resistance, potentially generating nutrient-mediated trade-offs across the nutritional landscape. Here, we used the GFN to examine how macronutrient balance structures growth, immune function, stress responses, and pathogen resistance in the specialist herbivore *Plutella xylostella* (Lepidoptera: Plutellidae), a major pest of economically important Brassicaceae crops worldwide (Talekar & Shelton, 1993). *P. xylostella* was the first insect pest documented to evolve field resistance to *Bacillus thuringiensis* (Bt) (Tabashnik *et al*., 1990), positioning it as a valuable model for exploring how nutritional state affects immune competence and survival under Bt exposure, with implications for both ecological understanding and resistance management. By manipulating dietary P:C ratios across a nutritional landscape, we quantified how variation in macronutrient composition drives multiple functionally linked traits. Specifically, we tested whether (i) growth is primarily constrained by protein availability, (ii) immune traits peak at intermediate P:C balance, and (iii) antioxidant and detoxification responses exhibit distinct nutrient-dependent patterns across the P:C landscape. We further assessed survival following *Bt* challenge to link nutrient-dependent physiological responses with pathogen resistance. By integrating life-history traits, immune parameters, oxidative responses, and infection outcomes within a unified nutritional framework, we tested whether a single macronutrient balance can simultaneously support multiple physiological functions or instead imposes competing physiological demands in this specialist herbivore.

## 2. Materials and methods

### 2.1 Insect rearing conditions

*Plutella xylostella* (Lepidoptera: Plutellidae) larvae were collected from cabbage fields in Nadia district, India, and maintained for several generations on a wheat germ-based basal artificial diet (F9441B, Frontier Agricultural Sciences) in the laboratory. Insect cultures were maintained in a growth chamber at 23±2 °C, 68±5% relative humidity, and a photoperiod of 16 h light: 8 h dark. Adult insects were provided with a 15% honey solution (v/v) diet dipped in cotton balls.

### 2.2 Chemicals

All substrates and chemicals were purchased from Sigma-Aldrich unless otherwise specified.

### 2.3 Dietary manipulation

The basal artificial diet was prepared according to the manufacturer’s protocol (F9441B, Frontier Agricultural Sciences). Per 100 mL distilled water, 16 g of a dry mix (acid casein, pasteurized wheat germ, stabilized sodium propionate, sorbic acid, and methyl paraben), 1.1 g agar, 1.0 g vitamin mix, 0.02 g chlortetracycline, and 700 µL linseed oil were combined and mixed thoroughly until homogeneous.

The basal diet’s protein and carbohydrate content was estimated as 22 g and 36 g per 100 g diet, respectively, using the Lowry method for protein and the phenol–sulfuric acid method for carbohydrate, with methodological modifications detailed in Supplementary Material S1. This composition served as the baseline for formulating experimental diets, to which casein and sucrose were added as supplementary protein and carbohydrate sources, respectively. Because the basal diet’s fixed protein and carbohydrate levels precluded full orthogonal manipulation of these nutrients, supplement levels were instead varied non-orthogonally to generate a dietary nutrient space for response surface analysis, following the GFN (Simpson & Raubenheimer, 2012). Diets were designed to span a range of P:C ratios from 1:1 to 1:2 (Table 1).

**Table 1:** Composition of experimental diets spanning the protein–carbohydrate nutritional landscape.

| Diet ID | Total protein (P)<br>(g /100g diet) | Total carbohydrate<br>(C)<br>(g /100g diet) | P: C |
| --- | --- | --- | --- |
| 1 | 22 | 36 | 1:1.6 |
| 2 | 24 | 36 | 1: 1.5 |
| 3 | 24 | 40 | 1:1.6 |
| 4 | 24 | 44 | 1:1.8 |
| 5 | 24 | 52 | 1:2 |
| 6 | 22 | 38 | 1:1.7 |
| 7 | 26 | 38 | 1:1.4 |
| 8 | 30 | 38 | 1:1.2 |
| 9 | 38 | 38 | 1:1 |

### 2.4 Larval rearing on defined diets and life-history measurements

Freshly hatched larvae were individually assigned to one of the experimental diets that varied in protein and carbohydrate content as described above (n = 10 larvae per diet). Larvae that completed development were transferred to pupation plates, while individuals exhibiting prolonged immobility and a melanized appearance were recorded as dead. Larval and pupal performance were quantified using the following metrics. Relative larval weight (RLW) was calculated as the difference between initial and final larval weights divided by the total number of days to pupation (Kharel *et al*., 2026). Pupal weight (PW) was recorded one day post-pupation, and pupal period (PP) was defined as the interval between pupation and adult emergence (Dhillon *et al*., 2021). All weight measurements were taken using a Mettler Toledo semi-microbalance (XPR105). Larval survival (LS) was expressed as the percentage of individuals reaching pupation relative to the initial cohort (Kharel *et al*., 2026).

### 2.5 Cellular immunity and melanisation assays

To quantify cellular immunity, hemolymph was collected from newly molted fourth-instar larvae reared on each experimental diet. Ten larvae were utilized for each diet treatment, with one larva assigned to each of 10 wells of Teflon-coated diagnostic slides (8 mm, 10 wells) (n = 10). Larvae were anesthetized on ice for 15 min, surface-sterilized with 70% ethanol, and hemolymph was extracted from proleg using sterile forceps. Samples were immediately mixed with ice-cold phosphate-buffer saline (PBS) in the slide well and fixed in 4% formaldehyde. After washing with PBT (0.3% Triton X-100 in PBS), samples were stained with Alexa Fluor 488 for 2 h in the dark and mounted with DAPI Fluoroshield. Hemocytes were visualized with an Olympus IX81 epifluorescence microscope. Total and differential hemocyte counts (THC and DHC) were quantified following literature protocol (Sasidharan *et al.,* 2025) and expressed as cells mm⁻².

For the phenoloxidase (PO) activity assay, hemolymph was collected from 20 newly molted fourth-instar larvae per dietary treatment. Hemolymph from two larvae was pooled, collected in 10 µL of ice-cold PBS, and transferred to a 96-well plate containing 30 µL of PBS. Subsequently, 10 µL of 10 mM L-DOPA was added, and samples were incubated in the dark for 1 h. Absorbance was measured at 490 nm using a BioTek Cytation 5 multimode reader (Sasidharan *et al.,* 2025).

### 2.6 Pathogen challenge and survival assay

Immune competence was assessed by quantifying larval survival following infection with the entomopathogenic bacterium Bt, (Deans *et al.,* 2016). *B. thuringiensis* var. *kurstaki* (Greenlife Biotech) was prepared at a concentration of 2 × 10□ CFU ml□¹. Larvae (n = 10) were infected by pricking the proleg with a needle dipped in the bacterial suspension (Behrens *et al*., 2014) and then returned to their respective diets, where they were maintained until death or pupation. Control larvae were pricked identically with a needle dipped in sterile 0.1 M PBS. Mortality was recorded at 24, 48, and 72 h post-infection (1 dpi, 2 dpi, 3 dpi).

### 2.7 Antioxidant and detoxification enzyme assays

Enzyme extracts were prepared following Shen *et al*. (2020), with modifications. Three independent biological replicates were established for each dietary condition, each comprising a pooled sample of ten fourth-instar *P. xylostella* larvae. Larvae were surface-sterilized and homogenized in 1 mL of ice-cold enzyme extraction buffer (50 mM sodium phosphate buffer in 10% glycerol, pH 7). Homogenates were centrifuged at 8000 rpm for 15 minutes at 4°C, and the supernatant was collected for enzyme assays. Extracts were stored at -20°C until analysis.

Superoxide Dismutase (SOD) activity was measured using the pyrogallol autoxidation method (Marklund & Marklund, 1974). The reaction mixture contained 100 μL of 100 mM potassium phosphate buffer, 1.5 mM of freshly prepared pyrogallol in 10 mM HCl and 50 μL of enzyme extract in a 96-well plate. Absorbance at 440 nm was recorded at 1 min intervals for 60 min. Enzyme activity was calculated from the rate of inhibition of pyrogallol autoxidation, with one unit defined as the amount of enzyme required to cause 50% inhibition, and expressed as U/mg protein. Protein concentration was determined using the Bradford assay (Bradford, 1976).

Glutathione-S-Transferase (GST) activity was quantified by measuring the conjugation of reduced glutathione (GSH) with 1-chloro-2,4-dinitrobenzene (Habig *et al*., 1974). The reaction mixture comprised 186 μL of 50 mM sodium phosphate buffer (pH 6.8), 2 μL of 100 mM GSH, 2 μL of enzyme extract, and 10 μL of 30 mM CDNB in a 96-well plate. The absorbance at 340 nm was recorded in 1-min intervals for 30 min, and enzyme activity was expressed as µmol mL⁻¹ 30 min⁻¹.

Carboxylesterase (CarE) activity was measured following van Asperen (1962), with modifications, using 1-naphthyl acetate as the substrate. 15 μL of the enzyme extract was incubated with 135 μL of the substrate at 4°C in the dark for 1 hour, followed by the addition of 50 μL of DBLB reagent (1% fast blue B salt and 5% SDS in a 40 mM phosphate buffer in a 2:5 ratio). After incubation at room temperature for 15 min, absorbance was measured at 590 nm. Enzyme activity was calculated using a 1-naphthol standard curve and expressed as nmol 1-naphthol.

Lipid peroxidation (LPO) was assessed using malondialdehyde (MDA) as a marker of oxidative damage (Siddique *et al*., 2012). Enzyme extract (100 μL) was mixed with 650 μL of 1-methyl-2-phenyl indole solution (0.064 g in 30 mL acetonitrile and 10 mL methanol), followed by 150 μL of 37% HCl. The mixture was incubated at 90°C for 45 min and then cooled on ice. After centrifugation, 200 μL of supernatant was transferred to a 96-well plate, and absorbance was measured at 600 nm. MDA concentration was determined using a 1,1,3,3-tetramethoxypropane standard curve and expressed as nmol MDA.

### 2.8 Statistical analysis

All analyses were performed in R v4.4.0 (R Core Team 2024). Effects of dietary protein and carbohydrate concentrations on trait responses were analyzed using second-order polynomial response surface models within the GFN framework. Models included linear terms for protein (P) and carbohydrate (C), their quadratic terms (P² and C²) and their interaction (P × C), allowing assessment of linear, non-linear and interactive effects on trait performance. Model assumptions were evaluated by a visual examination of residual diagnostic plots, and Shapiro–Wilk test used as a supplementary assessment of residual normality. Throughout the experiment, all larvae were housed individually, treating each larva as an independent experimental unit. Consequently, there was no nested or repeated-measures structure present in the study design. Traits with normally distributed residuals were analyzed using linear models (LMs), whereas non-normal responses were analyzed using generalized linear models (GLMs). Binomial responses were fitted using GLMs with a binomial error distribution and logit link function. The significance of model terms was assessed using Type II tests: F-tests for LMs and Type II likelihood-ratio tests for GLMs. Nutritional landscapes were visualized using thin-plate spline interpolation and contour plots implemented using the *fields* package in R (Nychka *et al*., 2021). For multivariate visualization, mean trait values for each diet were standardized as *Z*-scores before generating the diet × trait heatmap using the *ComplexHeatmap* package (Gu, 2022). Hierarchical clustering of diets and traits was performed using Euclidean distance and complete-linkage clustering. Statistical significance was determined at α = 0.05.

## 3. Results

### 3.1 Macronutrient balance differentially structures growth, development and survival

Larval growth varied predictably across experimental nutrient space, with the highest biomass occurring at intermediate to high protein concentrations and increasing carbohydrate availability (Figure 1a). Dietary protein had the strongest influence on larval weight (LRχ² = 36.52, *P < 0.001*), although dietary carbohydrate also contributed significantly (LRχ² = 12.05, *P < 0.001*). A significant quadratic effect of protein (LRχ² = 11.81, *P = 0.00059*) indicated a nonlinear response within the experimental nutrient space, whereas carbohydrate showed a predominantly linear effect. No P × C interaction was detected, indicating that the growth responses to protein and carbohydrate were independent rather than synergistic.

**Figure 1.**
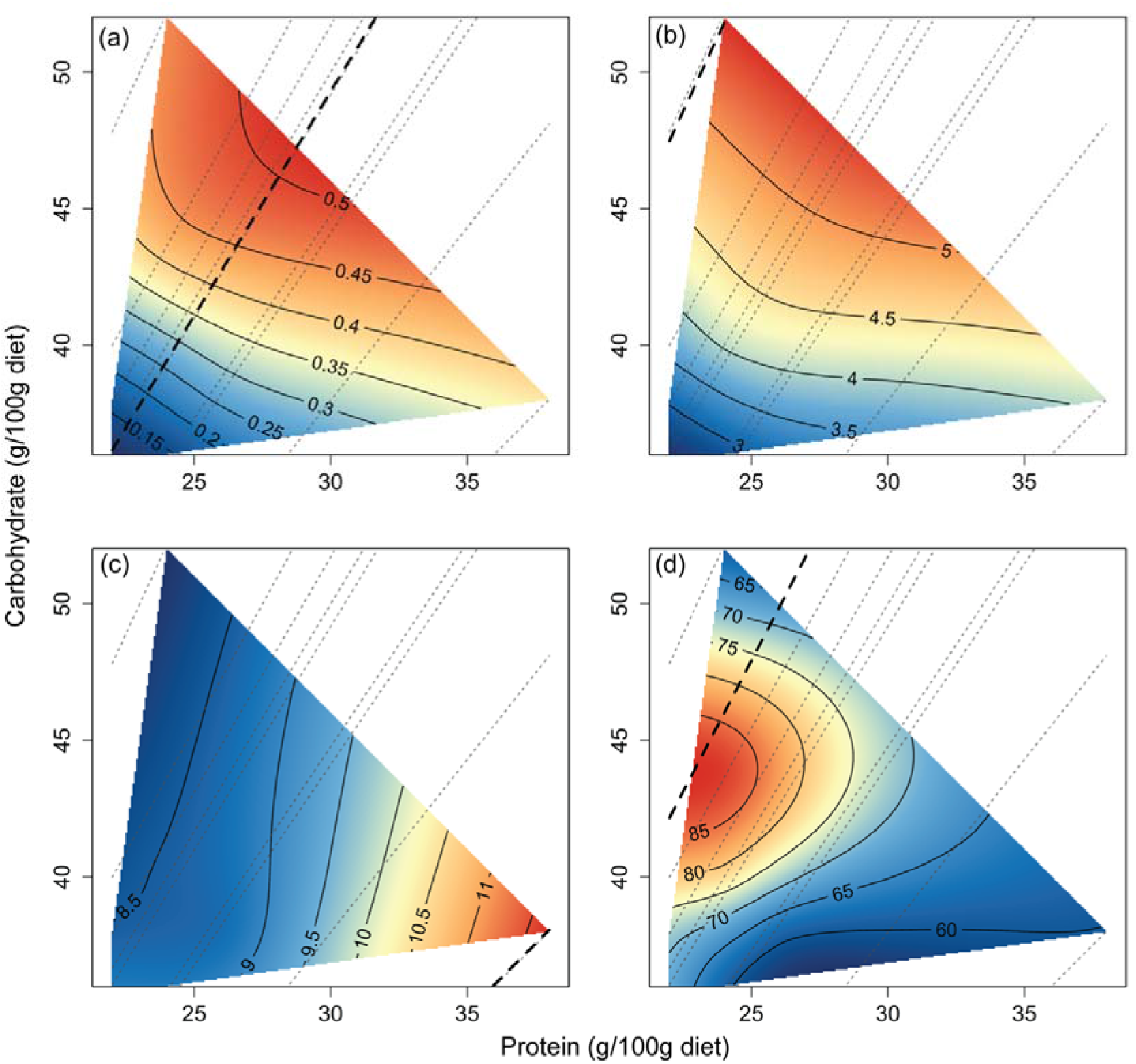
Performance landscapes of larval life-history traits in the nutritional space of *Plutella xylostella*. The two-dimensional space is defined by dietary protein (P) and carbohydrate (C) concentrations (g per 100 g diet). Trait responses were visualized using thin-plate spline interpolation, with color gradients indicating performance from low (blue) to high (orange). Gray dashed lines represent nutritional rails corresponding to fixed P: C ratios of the experimental diets, and the thick black dashed line denotes the ridge of maximal performance. Panels show (a) relative larval growth rate (mg day⁻¹), (b) pupal weight (mg), (c) pupal period (days), and (d) larval survival (%)

Pupal weight showed a response surface similar to larval growth, increasing with both dietary protein and carbohydrate availability, although protein exerted stronger influence on biomass (Figure 1b). In contrast, pupal period responded primarily to carbohydrate availability; larvae reared on protein-rich diets took longer to develop (∼11 days), whereas carbohydrate-rich diets developed faster (∼8.5 days). Pupal period was significantly influenced by carbohydrate (F = 5.66, *P* = *0.02*), consistent with a linear response surface (Figure 1c). The response surface indicated an apparent region of higher larval survival (∼85%) under relatively low-protein and intermediate-carbohydrate conditions (Figure 1d). However, the statistical results show neither dietary protein nor carbohydrate, nor their quadratic terms (P^2^ and C^2^), have effects on survival and no P × C interaction was detected across experimental diets (Supplementary material Table S1).

### 3.2 Macronutrient balance differentially regulates cellular immunity

Morphological analysis of *Plutella xylostella* larval hemocytes revealed three distinct hemocyte types: granulocytes, oenocytoids, and plasmatocytes, consistent with an earlier report (Ghosh *et al*., 2022). Total hemocyte counts (THC) varied primarily along the protein axis of the experimental nutrient space, with the highest values occurring at the intermediate dietary protein concentration (Figure 2a). Protein significantly influenced THC with a strong quadratic effect (F = 12.12, *P* < 0.001), whereas carbohydrate showed neither linear nor quadratic effects. No P × C interaction was detected, indicating that variation in hemocyte abundance was driven predominantly by protein availability.

**Figure 2.**
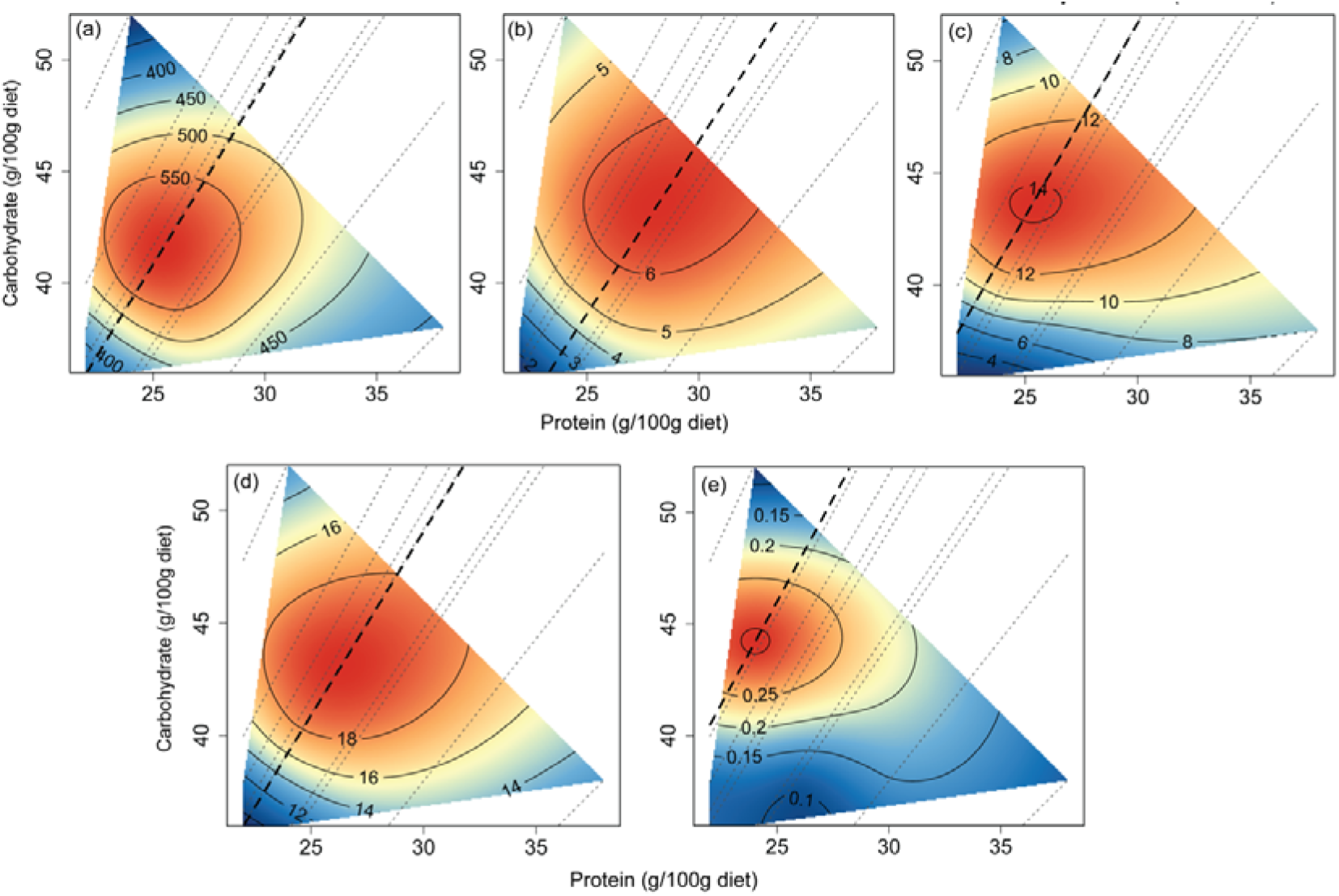
Nutritional performance landscapes for cellular and humoral immune traits of *Plutella xylostella*. The two-dimensional nutritional space is defined by protein (P) and carbohydrate (C) concentrations (g per 100 g diet). Performance surfaces were visualized using thin-plate spline interpolation, with red regions indicating higher immune responses and blue regions indicating lower responses. Gray dashed lines radiating from the origin represent the nutritional rails of the fixed-ratio diets, and the thick black dashed line denotes the ridge of maximal response across the nutrient space. Panels show (a) Total hemocyte counts (cells/mm^2^), (b) Plasmatocytes count (cells/mm^2^), (c) Oenocytoids count (cells/mm^2^), (d) Granulocytes count (cells/mm^2^) and (e) Phenoloxidase activity

The abundance of individual hemocyte types demonstrated distinct yet broadly consistent nutritional patterns across the experimental nutrient spectrum (Figure 2b–d). Plasmatocytes reached their peak abundance (>6 cells mm□²) on carbohydrate-rich diets with moderate protein (Figure 2b). Both protein (LRχ² = 27.45, *P < 0.001*) and carbohydrate (LRχ² = 17.22, *P < 0.001*) significantly influenced plasmatocyte abundance, with notable quadratic effects for protein (LRχ² = 20.61, *P < 0.001*) and carbohydrate (LRχ² = 12.69, *P < 0.001*), confirming a nonlinear relationship within the experimental nutrient space. Oenocytoids exhibited their highest abundance (∼14 cells mm□²) at intermediate levels of protein (Figure 2c). Protein had the strongest impact on oenocytoid abundance (LRχ² = 117.33, *P < 0.001*), accompanied by a marked quadratic response (LRχ² = 94.82, *P < 0.001*), whereas carbohydrate played a lesser but still significant role (LRχ² = 10.80, *P = 0.001*). Granulocyte abundance also reached approximately 19 cells mm⁻² under intermediate nutrient concentrations (Figure 2d). Protein (LRχ² = 43.13, *P < 0.001*) and carbohydrate (LRχ² = 9.81, *P = 0.0017*) both significantly influenced granulocyte abundance, with significant quadratic responses for both macronutrients (P^2^: LRχ² = 39.76, *P < 0.001*; C^2^: LRχ² = 11.21, *P < 0.001*). Across all hemocyte types, the absence of significant P × C interactions indicates that protein and carbohydrate contributed largely additive effects, while each hemocyte population demonstrated a distinct nutrient balance (Supplementary material Table S2).

Phenoloxidase (PO) activity mirrored these patterns, peaking at intermediate protein levels and covarying with oenocytoid abundance, consistent with their role as a primary enzyme source (Figure 2c, e). PO activity was driven primarily by protein, with a strong quadratic response (LRχ² = 26.21, *P* < *0.001*), while carbohydrate and the P × C interaction term was not significant (Figure 2e). Together, these results indicate that immune investment is structured by macronutrient balance, with protein imposing a primary constraint while distinct immune components exhibit trait-specific responses across the nutritional landscape.

### 3.3 Protein-dependent survival dynamics following *Bacillus thuringiensis* infection

The survival responses following infection with Bt showed distinct temporal patterns in the experimental nutritional space (Figure 3a–c). At 24 hours post-infection (1 dpi), survival remained high across nutrient combinations, peaking at intermediate protein and carbohydrate levels (Figure 3a), although statistical analysis indicated only a marginally non-significant quadratic trend for protein (LRχ2 = 3.44, *P=0.064*) and no significant linear effect of either macronutrient. By 48 hours post-infection (2 dpi), survival declined to approximately 75%, and mortality became more pronounced in nutritionally imbalanced diets (Figure 3b).

**Figure 3.**
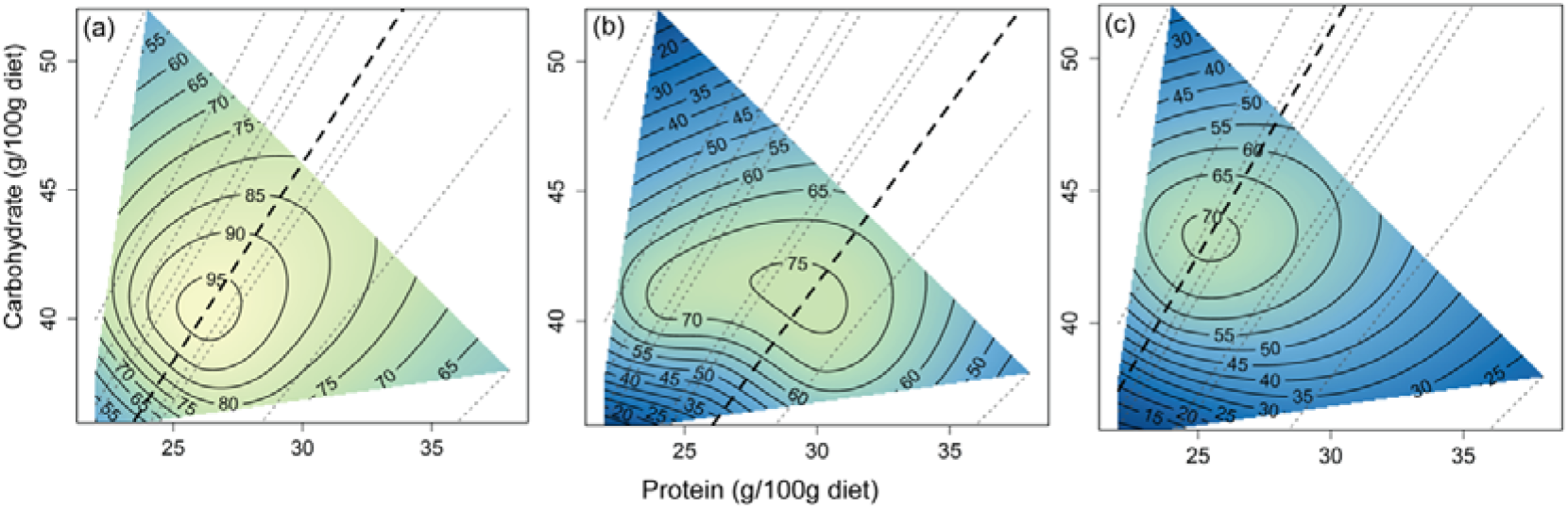
Nutritional performance landscapes for *Plutella xylostella* larval survival following *Bacillus thuringiensis* (Bt) infection at different days post-infection (dpi). The two-dimensional nutritional space is defined by protein (P) and carbohydrate (C) concentrations (g per 100g diet). Performance surfaces were visualized using thin-plate spline interpolation, with yellow regions indicating higher survival and blue regions indicating lower survival. Gray dashed lines radiating from the origin represent the nutritional rails of the fixed-ratio diets. The thick black dashed line denotes the ridge of maximum performance across the nutrient space. Panels show larval survival (%) at (a) 1 dpi, (b) 2 dpi, and (c) 3 dpi

Protein emerged as the principal factor influencing survival at this time point, with significant linear (LRχ² = 6.33, *P = 0.0119*), and quadratic (LRχ² = 9.59, *P = 0.002*) effect, indicating that survival was maximized at intermediate dietary protein levels. This trend intensified by 72 hours post-infection (3 dpi), where survival rates again peaked near 70%, at a P:C ratio of approximately 1:1.7 (Figure 3c). The effect of protein remained robust with significant linear (LRχ² = 12.24, *P < 0.001*) and quadratic (LRχ² = 11.93, *P < 0.001*) associations, while carbohydrate contributed minimally to survival at any time point. Together, these findings indicate that resistance to Bt infection is driven predominantly by protein nutrition, with intermediate protein levels conferring a substantial survival advantage - an effect that strengthened as the infection progressed (Supplementary material Table S3).

### 3.4 Nutrient-dependent divergence in antioxidant and detoxification responses

Antioxidant and detoxification enzymes displayed distinct nutritional response surfaces across the experimental nutrient spectrum (Figure 4a–d). SOD activity peaked at intermediate-to-high protein concentrations coupled with low carbohydrate availability (Figure 4a). Protein emerged as the primary determinant of SOD activity, with significant linear (F = 24.18, *P < 0.001*) and quadratic (F = 7.53, *P = 0.012*) effects. CarE activity showed a comparable response surface, with maximal activity occurring under similar nutritional conditions (Figure 4c). In contrast to SOD, CarE activity was significantly influenced by carbohydrate availability through both linear (F = 7.87, *P = 0.011*) and quadratic (F = 6.10, *P = 0.022*) effects, suggesting that detoxification capacity is more closely tied to carbohydrate than protein levels. Although the response surface showed relatively higher predicted GST activity in the low–protein/high–carbohydrate region of the nutritional spectrum, dietary protein was the only statistically significant predictor (F = 4.88, *P* = *0.038*), whereas carbohydrate and the interaction term were not significant.

**Figure 4.**
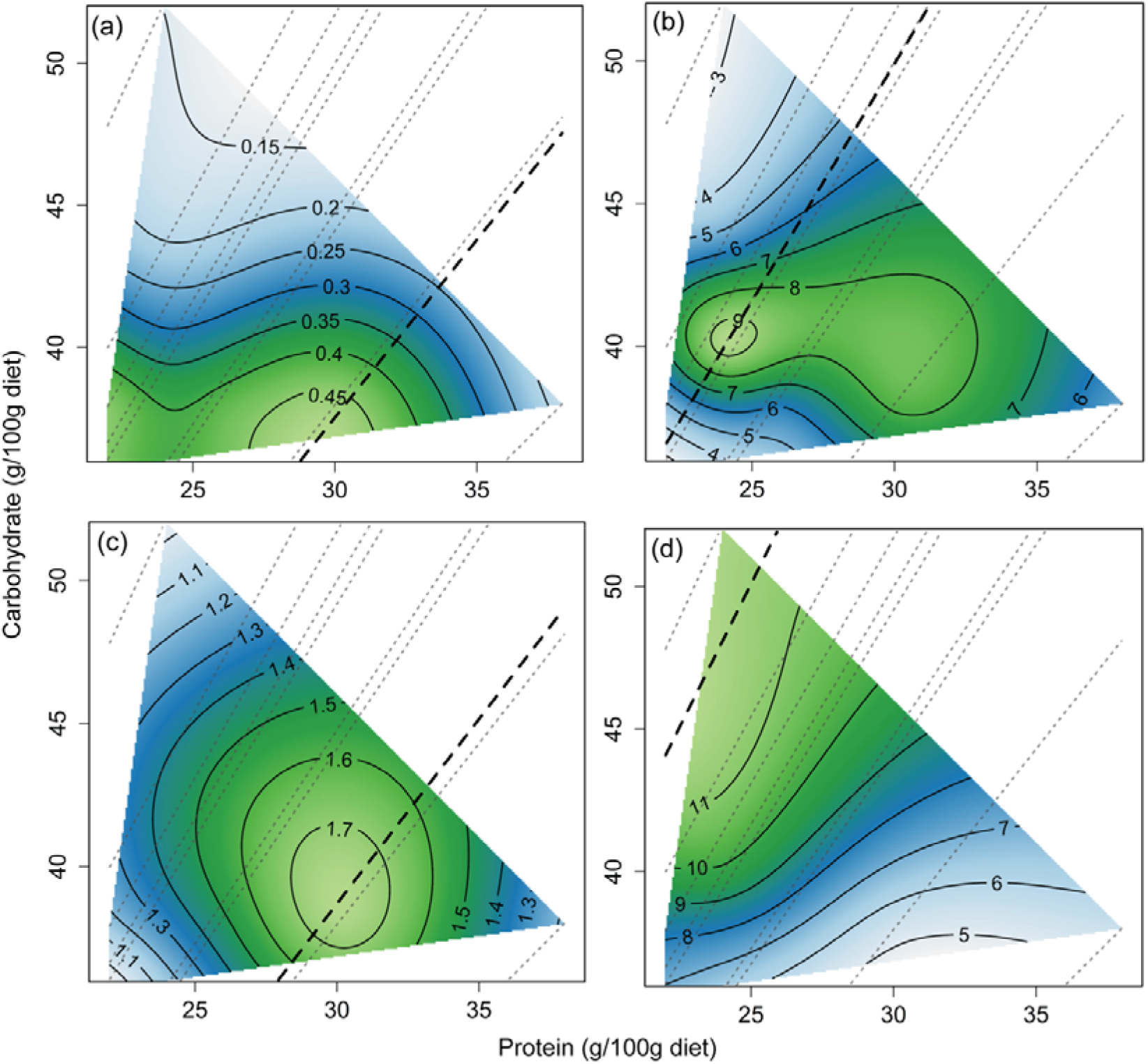
Performance landscapes of detoxifying enzymes and oxidative stress in the nutritional space of *Plutella xylostella*. The two-dimensional nutritional space is defined by dietary protein (P) and carbohydrate (C) concentration in the diet (g per 100g diet). Trait responses were visualized using thin-plate spline interpolation, with color gradients ranging from blue (low values) to green–yellow (high values). Gray dashed lines radiating from the origin represent nutritional rails corresponding to the fixed P: C ratios of the experimental diets, and the thick black dashed line denotes the ridge of maximal response across the nutrient space. Panels show (a) superoxide dismutase (SOD) activity (U mg⁻¹ protein), (b) lipid peroxidation (nmol MDA), (c) carboxylesterase (CarE) activity (nmol 1-naphthol), and (d) glutathione S-transferase (GST) activity (µmol mL⁻¹ 30 min⁻¹)

Lipid peroxidation, assessed by MDA content, was highest in protein-limited diets (Figure 4b), although neither macronutrient (P: LRχ^2^= 1.98, *P=0.159*; C: LRχ^2^ = 2.42, *P=0.119*) nor their interaction (LRχ^2^ =0.36, *P=0.546*) significantly predicted MDA content. Collectively, these findings illustrate that oxidative stress and detoxification responses are governed by distinct nutritional landscapes: SOD and CarE activity peak under similar surface conditions but are driven by different macronutrients, while GST activity and MDA accumulation follow contrasting nutritional patterns (Supplementary material S4).

### 3.5 Macronutrient composition structures two-clade divergence in immunocompetence, oxidative balance and fitness

Hierarchical clustering of row-standardized physiological and life-history traits revealed two distinct nutritional phenotypes across the nine experimental diets (see Figure 5). Diets clustered into two major groups according to their P:C balance, indicating that macronutrient composition was associated with coordinated shifts across multiple physiological traits rather than independent, diet-specific responses. The upper clade, comprising Diets 3, 4, 7, and 8, was characterized by positive Z-scores for oxidative stress, detoxification, and immune-related traits. Within this group, Diet 8 (P:C = 1:1.2) showed the highest standardized Z-scores for antioxidant and detoxification enzyme activities, along with high hemocyte abundance and survivability following a Bt challenge. Diet 7 (P:C = 1:1.4) clustered closely with Diet 8 but showed comparatively higher lipid peroxidation, indicating elevated oxidative stress despite an otherwise similar physiological profile.

**Figure 5.**
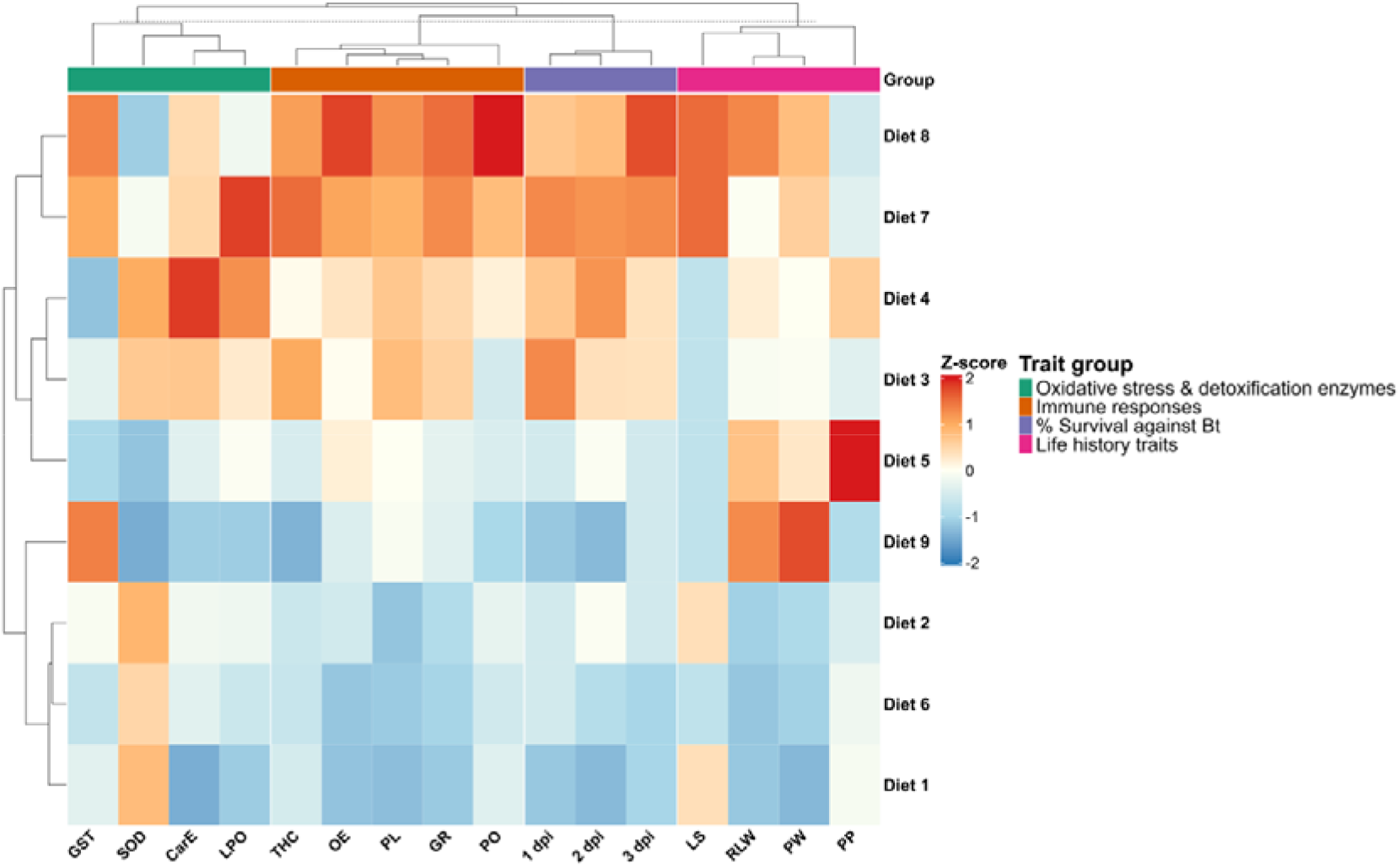
Hierarchical clustering heatmap showing the standardized phenotypic responses of *Plutella xylostella* larvae reared on nine artificial diets differing in protein and carbohydrate composition. Mean values for each trait within dietary treatments were standardized as Z-scores before hierarchical clustering to allow comparisons among variables on different scales. Warmer colors (red) indicate values above the mean for respective traits, whereas cooler colors (blue) indicate values below the mean. Both diets (rows) and traits (columns) were hierarchically clustered using Euclidean distance and complete linkage to identify patterns of similarity in dietary responses. Traits are grouped into four functional categories: oxidative stress and detoxification enzymes, immune responses, survival following *Bacillus thuringiensis* (Bt) infection, and life-history traits. The colored annotation bar above the heatmap denotes these trait categories. Abbreviations: GST: glutathione S-transferase; SOD: superoxide dismutase; CarE: carboxylesterase; LPO: lipid peroxidation; THC: total hemocyte count; OE: oenocytoids; PL: plasmatocytes; GR: granulocytes; PO: phenoloxidase; dpi: days post-Bt infection; LS: larval survival; RLW: relative larval weight; PW: pupal weight; PP: pupal period.

The second clade comprised Diets 1, 2, 5, 6, and 9, representing a contrasting physiological state characterized by generally lower Z-scores. These diets generally showed below–average or near–average responses across oxidative stress, detoxification, immune, and life–history traits. Diets 1, 2, and 6 consistently showed negative Z–scores across multiple trait categories. Diet 9, by contrast, presented a more heterogeneous profile, with below average standardized Z scores for oxidative stress and detoxification along with comparatively elevated values for select life–history traits, such as relative larval weight and pupal weight. Diet 5 showed predominantly below- to near-average standardized Z-scores across most physiological traits, with a notable exception of a markedly extended pupal period.

## Discussion

Our findings show that dietary macronutrient balance shapes growth, immunity, detoxification, and oxidative regulation in *Plutella xylostella* through largely distinct physiological responses. Integrating life-history traits, immunity, detoxification, and pathogen-resistance traits within a GFN framework revealed that these systems do not respond uniformly across the tested nutrient space; protein-associated and carbohydrate-associated processes occupy different regions of the landscape. This indicates that within the range of diets tested, no single P:C ratio maximizes all physiological functions simultaneously. While nutrient-dependent trait variation is documented across lepidopterans and other insects (Behmer, 2009; Cotter *et al*., 2011), our multidimensional approach shows how trait-specific responses generate this landscape in a specialist herbivore.

Growth responded additively to dietary protein and carbohydrate, with protein driving larval and pupal biomass – consistent with amino acid demand for structural protein synthesis and cuticle formation across molts (Rock & King, 1967; Muthukrishnan *et al*., 2020). The correlation between larval and pupal mass suggests larval nutrient acquisition carries over to metamorphic resources, when glycolytic and TCA-cycle activity are substantially reorganized (Hu *et al*., 2016; Zhou *et al*., 2015). Shorter developmental duration on carbohydrate-rich diets further points to carbohydrate as a key energy source during development.

Immune traits followed a nutritional pattern distinct from growth and one that maps onto the survival pattern we observed. Total hemocyte counts and oenocytoid densities were both highest on intermediate-protein diets, consistent with the nitrogen requirements of hemocyte production and, more specifically, of prophenoloxidase (PPO) synthesis by oenocytoids, the hemocyte subtype responsible for producing and storing PPO, the central precursor of melanization (Han *et al*., 1998; Lee *et al*., 2006; Wilson *et al*., 2019; González-Santoyo & Córdoba-Aguilar, 2012). Because Bt resistance in lepidopterans depends on the same protein-intensive machinery – antimicrobial peptides and serine-protease cascade that activates PPO (Kanost *et al*., 2004; Manniello *et al*., 2021), the diets associated with the highest total hemocyte and oenocytoid output were also those on which survival after Bt challenge was highest. This convergence is further supported by our clustering analysis, in which Diet 8 showed the highest standardized values for antioxidant and detoxification enzymes alongside high hemocyte abundance and survival following Bt challenge (Figure 5), directly linking protein-associated immune investment to pathogen resistance within a single dietary treatment. Together, this convergence, rather than the raw correlation between protein and survival alone, provides a mechanistic case that protein-driven melanization capacity - not just general protein sufficiency - plausibly contributes to the survival advantage. This parallels findings in *Spodoptera exempta*, where protein availability shaped antibacterial and PO activity (Povey *et al*., 2009).

Granulocytes and plasmatocytes, which contribute to phagocytosis and encapsulation showed different patterns: granulocyte abundance depended on both macronutrients, while plasmatocyte levels peaked on higher-carbohydrate, moderate-protein diet (Strand, 2026). This suggests that Bt resistance is not explained by protein-driven melanization alone, but reflects a composite of humoral and cellular defenses with somewhat divergent nutritional dependencies. Our data therefore support a correlational link between dietary protein and survival against Bt, pointing to hemocyte proliferation and in particular, oenocytoid/PPO-mediated melanization, rather than cellular immunity generally, as the likely mechanistic contributor.

The carbohydrate-associated processes were reflected in antioxidant and detoxification enzymes. SOD activity tracked protein availability, paralleling immune traits; CarE activity peaked under similar nutritional conditions but, unlike SOD, it was better explained by carbohydrate availability, suggesting its regulation may be more tied to energetic than protein status. GST activity was highest on carbohydrate-rich, protein-poor diets, consistent with its role in conjugating electrophiles and buffering oxidative stress (Felton & Summers, 1995). Similar diet dependent modulation of GST activity has also been reported in *Drosophila* (Strilbytska *et al*., 2022). MDA was nominally highest under protein-limited conditions but showed only weak overall dependence on diet, suggesting redox homeostasis was largely maintained across the tested range.

Hierarchical clustering across all traits reinforced this P:C partitioning: growth, oenocytoid-mediated melanization, SOD activity, humoral immune responses, and Bt resistance clustered together as protein associated traits while plasmatocyte abundance, CarE/GST activity, and developmental time clustered as carbohydrate associated traits. This matches a core GFN prediction – that performance reflects simultaneous resolution of multiple, sometimes competing, nutrient requirements, and generating trade-offs among competing nutrient requirements (Raubenheimer & Simpson, 1997; Wilson *et al*., 2019).

This partitioning is likely especially consequential for *P. xylostella*, an obligate Brassicaceae specialist that routinely encounters glucosinolate-derived allelochemicals (Textor & Gershenzon, 2008). Even with counter-adaptations such as glucosinolate sulfatases (Chen *et al*., 2023), sustaining detoxification, antioxidant defense, and immune function on chemically defended hosts imposes substantial metabolic cost (Futuyma & Moreno, 1988; Xia *et al*., 2017). Because our diets were glucosinolate-free artificial diets, the responses measured here represent the baseline demands; the added burden of host-plant chemistry remains an open question.

Collectively, macronutrient balance shapes growth, immunity, detoxification, and oxidative regulation through distinct, only partially overlapping responses, even across relatively narrow P:C gradients tested here. While most physiological functions diverged across this nutritional landscape, reflecting trade-offs among competing demands, protein availability emerged as a shared modulator linking immunity and resistance to Bt.

## Acknowledgments

Financial support from the Department of Biotechnology (DBT) and the Science and Engineering Research Board (SERB) POWER grant to RV is acknowledged. AG thanks Council of Scientific and Industrial Research (CSIR) for the fellowship. The authors thank the bioanalytical and imaging facilities at IISER Kolkata.

