## Supplementary material for "Macronutrient balance differentially regulates growth, immune function, and oxidative stress responses in *Plutella xylostella*"

**S1. Estimation of total protein and carbohydrate content of basal artificial diet**

The total protein content of the basal diet was estimated using a modified methodology based on Lowry et al. (1951). 100 mg of the sample was thoroughly homogenized with 10 mL of a 4% (w/v) sucrose solution, centrifuged at 4000 rpm for 6 min and the supernatant was stored at 4°C until further analysis. Bovine serum albumin (BSA) standards (0 to 2400 µg per reaction) and 1 mL of standard was mixed with 4 mL of alkaline copper reagent and incubated for 10 minutes. Then, 0.5 mL of Folin–Ciocalteu reagent was added to this mixture, incubated for an additional 30 minutes at room temperature, and absorbance was measured at 660 nm.

The total carbohydrate content of the basal diet was determined using the phenol-sulfuric acid method (DuBois et al., 1956), with some modifications. 100 mg of the basal diet was hydrolyzed with 5 mL of 2.5 N HCl in a hot-water bath for 3 h. After cooling the hydrolysate to room temperature, it was gradually neutralized with sodium carbonate until effervescence ceased. After adjusting the volume to 10 mL using Milli-Q water, the mixture was centrifuged at 4000 rpm for 6 min, and the supernatant was stored at 4°C until analysis. Glucose standard (0 to 1000 µg per reaction) was mixed with 1 mL 5% (w/v) phenol and 5 mL conc. H₂SO₄, adjusted to a final volume of 7 mL with Milli-Q water and then incubated in a water bath at 90°C for 10 min. After cooling, the absorbance was measured at 490 nm.

**Table S1: Effects of protein (P) and carbohydrate (C) on life history parameters.** Prior to analysis, the protein and carbohydrate values were mean-centred. The terms P^2^ and C^2^ represent the quadratic components. *P*-values were obtained using type II tests for LMs, and type II likelihood ratio tests for GLMs. F was used for LM (pupal period) and LR *χ^2^* for GLM (all other traits).

| Model terms | Relative larval weight (mg/day) | | Pupal weight (mg) | | Pupal period (days) | | Percent larval survival (%) | |
| --- | --- | --- | --- | --- | --- | --- | --- | --- |
|  | *χ^2^/F* | *P* | *χ^2^/F* | *P* | *χ^2^/F* | *P* | *χ^2^/F* | *P* |
| P | 36.52 | <0.001 | 57.63 | <0.001 | 1.72 | 0.193 | 2.44 | 0.118 |
| C | 12.05 | 0.001 | 14 | <0.001 | 5.66 | 0.02 | 1.19 | 0.276 |
| P^2^ | 11.81 | <0.001 | 9.83 | 0.0017 | 0.00039 | 0.984 | 3.01 | 0.083 |
| C^2^ | 3.6 | 0.058 | 3.5 | 0.061 | 2.21 | 0.141 | 0.18 | 0.668 |
| P × C | 1.63 | 0.2 | 0.004 | 0.951 | 0.51 | 0.479 | 0.23 | 0.635 |

**Table S2: Effects of protein (P) and carbohydrate (C) on immunity parameters.** Prior to analysis, the protein and carbohydrate values were mean-centred. The terms P^2^ and C^2^ represent the quadratic components. *P*-values were obtained using type II tests for LMs, and type II likelihood ratio tests were utilized for GLMs. F was used for LM (THC) and LR *χ^2^* for GLM (all other traits).

| Model terms | THC  (cells/mm^2^) | | Plasmatocyte count (cells/mm^2^) | | Oenocytoid  count (cells/mm^2^) | | Granulocyte count (cells/mm^2^) | | PO activity at 490 nm | |
| --- | --- | --- | --- | --- | --- | --- | --- | --- | --- | --- |
|  | *χ^2^/F* | *P* | *χ^2^/F* | *P* | *χ^2^/F* | *P* | *χ^2^/F* | *P* | *χ^2^/F* | *P* |
| P | 6.39 | 0.013 | 27.45 | <0.001 | 117.33 | <0.001 | 43.13 | <0.001 | 16.73 | <0.001 |
| C | 0.48 | 0.491 | 17.22 | <0.001 | 10.8 | 0.001 | 9.81 | 0.0017 | 0.23 | 0.629 |
| P^2^ | 12.12 | <0.001 | 20.61 | <0.001 | 94.82 | <0.001 | 39.76 | <0.001 | 26.21 | <0.001 |
| C^2^ | 2.05 | 0.156 | 12.69 | <0.001 | 4.3 | 0.038 | 11.21 | <0.001 | 0.36 | 0.547 |
| P × C | 0.47 | 0.497 | 0.27 | 0.601 | 0.01 | 0.924 | 0.65 | 0.421 | 1.32 | 0.25 |

**Table 3: Effects of protein (P) and carbohydrate (C) on larval survivability post-Bt pathogenesis.** Prior to analysis, the protein and carbohydrate values were mean-centred. The terms P^2^ and C^2^ represent the quadratic components. *P*-values were obtained using type II likelihood ratio tests for GLMs.

| Model terms | Percent larval survival post 24 hours of Bt infection | | Percent larval survival post 48 hours of Bt infection | | Percent larval survival post 2 72 hours of Bt infection | |
| --- | --- | --- | --- | --- | --- | --- |
|  | *χ^2^* | *P* | *χ^2^* | *P* | *χ^2^* | *P* |
| P | 1.98 | *0.159* | 6.33 | *0.0119* | 12.24 | *<0.001* |
| C | 2 | *0.157* | 3.06 | *0.08* | 1.03 | *0.311* |
| P^2^ | 3.44 | *0.064* | 9.59 | 0.002 | 11.93 | *<0.001* |
| C^2^ | 2.64 | *0.104* | 2.11 | 0.146 | 1.77 | *0.183* |
| P × C | 0.005 | *0.946* | 0.3 | 0.587 | 0.07 | *0.797* |

**T4: Effects of protein (P) and carbohydrate (C) on oxidative stress and detoxification.** Prior to analysis, the protein and carbohydrate values were mean-centred. The terms P^2^ and C^2^ represent the quadratic components. *P*-values were obtained using type II tests for LMs, and type II likelihood ratio tests were utilized for GLMs. F was used for LM (SOD and CarE activities) and LR *χ^2^* for GLM (other traits).

| Model terms | SOD activity  (U/mg protein) | | Lipid peroxidation  (nmoles of MDA) | | CarE activity  (nmoles of 1-naphthol) | | GST activity  (μmole/ml/30 min) | |
| --- | --- | --- | --- | --- | --- | --- | --- | --- |
|  | *χ^2^/F* | *P* | *χ^2^/F* | *P* | *χ^2^/F* | *P* | *χ^2^/F* | *P* |
| P | 24.18 | <0.001 | 1.983 | *0.159* | 1.271 | 0.272 | 4.885 | 0.038 |
| C | 0.72 | 0.406 | 2.427 | *0.119* | 7.868 | 0.011 | 1.559 | 0.226 |
| P^2^ | 2.36 | 0.139 | 3.15 | *0.076* | 2.458 | 0.132 | 1.474 | 0.238 |
| C^2^ | 7.53 | 0.012 | 1.29 | *0.256* | 6.096 | 0.022 | 0.3 | 0.59 |
| P × C | 0.0003 | 0.985 | 0.364 | *0.546* | 0.236 | 0.632 | 0.014 | 0.907 |
